# *In silico* analysis reveals the structural basis of TomEP specificity, a tomato extensin peroxidase

**DOI:** 10.64898/2026.03.10.710923

**Authors:** Zawar Hussain, Sumit Sharma, Ahmed Faik, Michael A. Held

**Affiliations:** Environmental & Plant Biology Department, Ohio University, Athens, OH 45701, USA; Department of Chemical and Biomolecular Engineering, Ohio University, Athens, OH 45701, USA; Department of Chemistry and Biochemistry, Ohio University, Athens, OH 45701, USA; Molecular and Cellular Biology Program, Ohio University, Athens, OH 45701, USA

**Keywords:** Cell wall, extensin, EPs, 3D structure, docking, Molecular Dynamics Simulation

## Abstract

**Background:** Extensin peroxidases (EPs) are class III plant peroxidases and are responsible for intermolecular covalent crosslinking of extensin (EXT) monomers to create scaffolds within plant cell walls. The formation of these scaffolds impacts plant development, mechanical wounding, and response to pathogen attacks. Therefore, elucidating the molecular mechanism controlling covalent crosslinking of EXT monomers is crucial for understanding cell wall deposition and potentially improving plant growth and adaptation. The focus of this work is to use *in silico* analysis to determine the structural characteristics of an EP from tomato (TomEP) to elucidate its specificity for crosslinking of EXT monomers.

**Results:** In this study the two-dimensional (2D) and three-dimensional (3D) structures of TomEP were determined using several advanced bioinformatics tools and compared to two other peroxidases: *Gv*EP1 (a known EP) and HRP-C (having a low affinity for EXT substrates). The results revealed that TomEP is a stable and hydrophilic protein with high thermal stability. The heme binding pockets of TomEP and *Gv*EP1 have more hydrophobic residues and larger volume and pocket area compared to HRP-C. Molecular docking at the active site, which includes a heme heteroatom, showed that the ligands consisting of the hydrophobic Tyrosine-X-Tyrosine [-Y-X-Y-] motifs (i.e., [-Y-K-Y-], [-Y-V-Y-], and [-Y-Y-Y-] found in EXTs, and their derivatives, Isodityrosine (IDT), Pulcherosine (Pul), Di-Isodytirosine (diIDT), bind perfectly to the active site of TomEP with dominant interactions of Val54, Ser94, Ala96 and Phe196 residues. Pulcherosine had the highest binding affinity, while [-Y-K-Y-] showed the lowest binding affinity. Molecular dynamics simulations showed that [-Y-X-Y-] motifs (and the derivative substrate ligands) remain bound to the active site of TomEP throughout the 100 ns long simulation. Furthermore, the binding of these substrates stabilized the protein structure.

**Conclusion:** These results may explain why TomEP is particularly well-suited for EXT crosslinking and will have significant implications on biochemistry, biotechnology, and the potential use of these EPs in crops improvement.

## Introduction

Most plant cells grow anisotropically, which is directed primarily by the yielding of the cell wall at precise locations in response to passive, constant turgor pressure [1]. It has been established that anisotropic cell elongation is dictated by a circumferential deposition of parallel cellulose microfibrils [2, 3]. However, cellulose microfibril deposition is not the only mechanism that regulates anisotropic cell elongation. EXT crosslinking may play a role in maintaining the assembly of the cell wall, which is critical for plant growth and development [4]. Plant cell walls contain a large variety of structural glycoproteins, which can constitute as much as 10% of the cell wall biomass [5]. Based on their amino acid composition, these glycoproteins are classified into six major groups, namely glycine-rich proteins, solanaceous lectins, EXTs, proline-rich proteins, proline/hydroxyproline-rich glycoproteins and arabinogalactan proteins (AGPs) [6]. Structural proteins can be covalently crosslinked into cell wall network and can affect wall mechanical properties and assembly. Among all the structural proteins, branched AGPs and the classical EXTs play important roles [7]. EXTs are found abundantly in the cell walls of Bryophytes, and plants [8] and help in the regulation of cell wall properties/assembly, plant development [4] and resistance against biotic and abiotic stresses [9]. Specifically, EXTs are involved in cell division [10, 11], seed germination [12], plant reproduction [13], disease and pathogen resistance [14] and environmental stress responses [15]. Despite this importance, our understanding of their crosslinking mechanism remains limited.

By definition, EXTs are characterized by the presence of [-Ser-Hyp_3-5_-]_n_ peptide repeats that are O-glycosylated with monogalactosyl (Gal) residues and short (3-5 residues) arabinofuranosyl (Ara*f*) oligosaccharide side chains on Ser and Hyp side chains respectively (Fig. 1A) making them rather hydrophilic [11, 16, 17]. These sugar decorations are responsible for the stabilization of EXTs’ rod-like structures [18] and facilitate intermolecular self-assembly of the crosslinked EXT network [19, 20, 21]. The hydrophilic repeats in EXTs are spaced by relatively hydrophobic [-Tyr-X-Tyr-] motifs, where X is typically (Tyrosine (Tyr, Y), Valine (Val, V), Serine (Ser, S), or Lysine (Lys, K)). The tyrosines in this motif can undergo *intra*molecular crosslinking to form isodityrosine (IDT) [22]. Free Tyr residues and IDT can undergo further covalent *inter*molecular crosslinking which is catalyzed by cell wall enzymes called EXT peroxidases (EPs) [16, 23]. Covalent crosslinking occurs to lock in place the self-assembled scaffold of EXT monomers through the formation of Pulcherosine (Pul), between an IDT and Tyr residues, and Di-IDT, between two IDT motifs (Fig.1A; Table.S1 (Supplementary Information) [9, 16].

**Fig.1.**
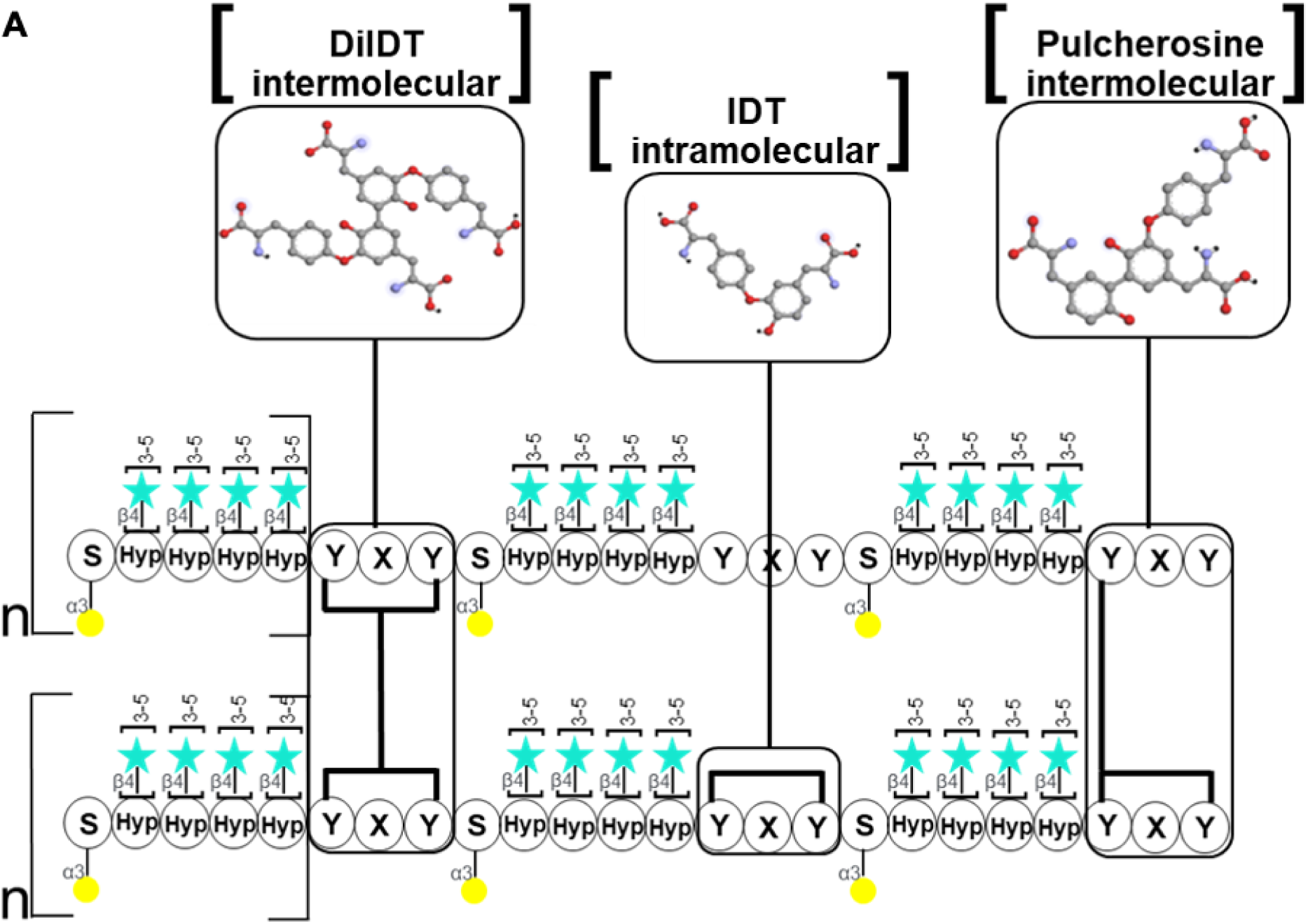
Extensins protein [Ser-Hyp3-5] motifs and glycosylation. **(A)** Schematic diagram of characteristic extensin [Ser-Hyp_3-5_] motifs and glycosylations, with EXT precursors self-aligned through amphiphilic interactions. The yellow circles represent galactopyranose and the light blue stars represent short (3-5 residues) oligoarabinofuranosyl side chains.

EPs are class III plant peroxidases and have been identified and characterized in several plant species including tomato [24], grapevine [25], lupin [26], and french bean [27]. Five promising EP candidates have been identified in Arabidopsis (*Arabidopsis thaliana*) for EXT crosslinking *in vivo* [28, 29], but their activity and kinetics for EXT crosslinking have not yet been confirmed *in vitro*. Of all the reported EPs, only TomEP has been shown to crosslink EXTs *in vitro* at physiologically relevant conditions (i.e., physiological amounts of enzyme, peroxide and EXT co-substrates) [24, 30]. In fact, TomEP is the only known EP that shows high crosslinking activity toward EXT monomers using very small amounts (∼5 ng) of enzyme [24]. TomEP was also found to be active on synthetic EXT analogues [23]. The amino acid composition of deglycosylated, TomEP-crosslinked EXTs revealed the presence of cross-linked products di-IDT as an intermolecular crosslinking between P3-type EXT monomers ([Ser-Hyp-Hyp-Hyp-Hyp-Ser-Hyp-Ser-Hyp-Hyp-Hyp-Hyp-Tyr-Tyr-Tyr-Lys]_n_) [23]. Later, Dong et al., (2015) identified the TomEP candidate gene (Solyc02g094180) using both proteomics and bioinformatics analyses and confirmed TomEP’s identity (aka CG5) and crosslinking activity for both P1 ([Ser-Hyp-Hyp-Hyp-Hyp-Val-Lys-Pro-Tyr-His-Pro-Thr-Hyp-Val-Tyr-Lys]_n_) and P3 ([Ser-Hyp-Hyp-Hyp-Hyp-Ser-Hyp-Ser-Hyp-Hyp-Hyp-Hyp-Tyr-Tyr-Tyr-Lys]_n_) type EXTs [30]. More recently, Mishler-Elmore et al., (2020) characterized the *in vivo* expression of TomEP [31]. Despite these advances, we still lack a mechanistic understanding of EXT crosslinking by EPs and what makes these enzymes specific to EXTs. One strategy to gain this information is to determine the 3D structure of TomEP and compared to other peroxidases.

Considering the difficulties in experimentally determining the 3D structure of proteins, *in silico* protein characterization provides a valuable alternative for determination of structural, dynamics, and biochemical characteristics. The 3D structure of many plant peroxidases have been predicted *in silico* through advanced bioinformatics tools such as peroxidases from rice [32], papaya [33], *Coprinus cinereus* [34], Arabidopsis [35], *Luffa acutangula* [36], and *Trachycarpus fortune* [37]. In this study, the 3D structure, binding pocket, and interacting surfaces between TomEP and the substrate ligands were predicted using AlphaFold program. PyMOL and ChimeraX were used for visualization. By searching Dali, a protein structure database searching server, closest homologous proteins based on 3D structural similarities were identified. CASTp program was used to measure the volume and area of TomEP’s catalytic cavity. Avogadro program was used to test the binding fitness of the substrate ligands ([-Y-X-Y-] and derivatives). PyRx and AutoDock Vina were used for molecular docking, Discovery Studio was used for docked complexes interactions analysis, while Maestro and Desmond programs were used for Molecular Dynamics Simulation (MDS) [38, 39]. As expected, our results showed that TomEP has a 3D structure that is similar to known peroxidases from class III. However, the hydrophobic residues surrounding the active cavity and the volume and area of binding pocket were specific to TomEP and *Gv*EP1, which may explain the affinity towards EXTs. Furthermore, molecular docking and MDS showed that EXT substrate motifs [-Y-X-Y-] (i.e., [-Y-K-Y-], [-Y-V-Y-], and [-Y-Y-Y-] and the derivatives (i.e., IDT, Pul, Di-IDT) (Table.S1) bind favorably to the TomEP active site and remain stable. We also identified some amino acid residues in the binding cavity that are engaged with all of the substrate ligands we examined, suggesting their potential roles in catalysis. These findings highlight TomEP’s structural suitability for EXT crosslinking.

## Methods

### Assessment of TomEP’s physicochemical properties

Physiochemical properties of TomEP were predicted using the ProtParam tool of ExPASy database (https://web.expasy.org/protparam/) [40]. This server provides molecular weight, amino acid composition, molecular formula, theoretical isoelectric point (pI), total number of positive (Arg + Lys) and negative (Asp + Glu) amino acid residues (+R/-R), Extinction Coefficient (EC), Instability index (II), and the grand average of hydropathicity (GRAVY) [35].

### Secondary structure and sequence annotation of TomEP

Secondary structure of TomEP was predicted using the secondary structure prediction tools namely, PSI-blast-based secondary structure prediction (PSIPRED) (Web: http://bioinf.cs.ucl.ac.uk/psipred/) and the self-optimized prediction method with alignment (SOPMA)(URL:https://npsaprabi.ibcp.fr/cgibin/npsa_automat.pl?page=/NPSA/npsa_sopma.html [41–43]. The Sequence Annotated by Structure (SAS) server from EMBL-EBI (URL: https://www.ebi.ac.uk/thornton-srv/databases/sas/) was used to confirm the TomEP secondary structure and define its active and binding sites. The active and binding sites were further confirmed from UniProt (https://www.uniprot.org/uniprotkb/A0A3Q7FC86/entry) and COFACTOR ([COFACTOR server](https://seq2fun.dcmb.med.umich.edu/COFACTOR2/)) server. The binding pocket was then defined and compared with Horse Radish Peroxidase (HRP)-C and grapevine extensin peroxidase (*Gv*EP1) using CASTp server (http://sts.bioe.uic.edu/castp/index.html?3igg).

### 3D structure prediction and validation

The 3D structure of TomEP was predicted using AlphaFold 3, developed by DeepMind and EMBL-EBI (https://alphafoldserver.com/welcome/). To check the accuracy and reliability of the Alphafold 3D structure, the predicted 3D structure was assessed and evaluated using Ramachandran plot, MolProbity, Errat, ProSA, PROCHECK and ProQ servers. Furthermore, we identified the closest structural homologous models available in the Protein Data Bank using the DALI server (http://ekhidna2.biocenter.helsinki.fi/dali/). DALI identified top homologous crystal structures based on z-scores and superimposed the first homologous structure and two other reference peroxidases (Horseradish peroxidase (HRP-C) and Grapevine peroxidase (*Gv*EP1: extensin peroxidases) with TomEP. To check the structural stability, the 3D structure was subjected to isothermal isobaric ensemble molecular dynamics simulation for 100 ns at 300 K and 1 atmosphere pressure.

### Molecular docking and enzymes-substrate ligands interaction

Molecular docking of substrate ligands on TomEP’s active site was carried out to determine the binding affinity, analyze the types of interactions, and identify key residues involved in the enzyme-ligand catalysis. Six substrate ligands were prepared using PyMOL and Avogadro software. The substrate ligands and their minimum energy 3D structures are listed in Table S1. The substrates and TomEP’s 3D structures were imported to the PyRx autodock tool and their energy was minimized. Among the substrates, three were [-Y-X-Y-] substrate ligands (i.e. [-Y-V-Y-], [-Y-K-Y-] and [-Y-Y-Y-]) and three were crosslinked derivatives (i.e. IDT, Pul and DiIDT). The substrate ligands were docked onto TomEP using Vina Wizard tool in PyRx. First, a blind docking over the whole protein surface was done, followed by a more selective one, reducing the search space to a box centered over the most frequent binding site found in the previous run. For each ligand, the docked complexes with the lowest binding affinity were analyzed using Discovery Studio, PyMOL and Chimera X.

### Molecular dynamics simulations

Molecular dynamics simulations of all the docked complexes were performed using the Desmond package (Schrodinger v05.1/2023.2) (Maestro v11.9 and Desmond v5.7) utilizing the Ohio State Supercomputer Center [44], to check stability and catalytic interactions [45]. The interactions were modeled using the Optimized Potentials for Liquid Simulations (OPLS3) force field [46, 47]. The protein complex was solvated by water. Water was modeled using the simple point charge enhanced (SPC/E) model. Cubic simulation box was set with 10 Å × 10 Å × 10 Å from the nearest protein atom. The system was made charge-neutral by adding Na^+^ ions [48]. The energy of the system was minimized, and isothermal isobaric (NPT) ensemble simulations were performed for 100 ns. The temperature was maintained at 300 K and the pressure at 1 atm using the Berendsen thermostat and barostat [49]. Coulombic interactions were calculated using particle mesh Ewald method with a cut-off distance of 9 Å for the real space part. Water molecules were kept rigid using the SHAKE algorithm [50]. The 100 ns trajectory of the protein-ligand complex was analyzed to determine if the ligand remained bound to the protein’s binding site.

## Results

### TomEP is a stable and hydrophilic protein with high thermal stability

TomEP has a molecular weight of 35.39 kDa and a theoretical pI value of 4.6, suggesting that the protein is slightly acidic or negatively charged at pH 7. The protein contains 39 negatively charged residues (Asp and Glu), 28 positively charged residues (Arg and Lys), and does not contain Tryptophan (Trp). The instability index (II) is one indicator of stability of a protein, with values below 40 suggesting that the protein is stable [51]. TomEP has an II equal to 29.54 indicating that the protein is stable. The aliphatic index (AI) is a measure of the volume of protein comprising of aliphatic side chains. Since protein folding is known to be dictated by the hydrophobic effect, a large AI suggests higher thermal stability of protein, although this is a coarse measure [52]. The AI of TomEP is 85.82, which indicates high thermal stability. The GRAVY (Grand Average of Hydropathicity) of TomEP was found to be -0.02, suggesting that the protein is hydrophilic (Table.S2, Fig.S1).

TomEP’s secondary structure was determined using the PSIPRED and SOPMA tools. SOPMA showed that TomEP consists of 40.62% alpha-helix, 16.92% extended strand, 5.23% beta strands and 37.23% random coils (Table S3). PSIPRED analysis revealed that the secondary structure contains helices, strands and coils shown in amino acids sequence form with small letter legends (Fig. S3A). Sequence Annotated by Structure (SAS) analysis is useful for identifying binding sites on the protein [53, 54]. SAS analysis of TomEP allowed the annotation of various structural parts of its secondary structure, including sheets, beta strands, helices, residue contacts to metals, catalytic residues, and active sites (Fig. S3B).

### TomEP has a 3D structure with high similarity to other class III peroxidases

TomEP’s 3D structure was generated using AlphaFold 3 (Fig. 2A). The predicted structure was compared to the closest, structurally homologous crystal structures available in the Protein Data Bank (PDB) identified through DALI search [55]. The search identified several peroxidases, and the top 10 were selected for further analysis (Table S4). All the identified homologs were also class III peroxidases. Among these homologs, we selected two peroxidases, HRP-C and *Gv*EP1 for comparison with TomEP. TomEP structure superimposed closely to the structures of these two peroxidases with the Root Mean Squared Deviation (RMSD) value of 0.838 Å which indicates high similarity at the 3D structure levels (Fig. 3A). RMSD is calculated by aligning the protein backbones of the structures and then finding the root mean squared distance between the heavy atoms of the two structures. Next, molecular dynamics simulations were used to determine the structural stability at constant temperature and pressure of the predicted 3D structure of TomEP for 100 ns. Results from the simulation showed that the RMSD values remained stable at around 1.8 Å with the largest deviation of 2.4 Å. In addition, the root mean square fluctuations (RMSF) values of all residues were within 2.4 Å (Fig. S2C and S2D). Panel (A) and (B) of Fig. S2 show the beginning and the last configuration from the MDS, indicating that the protein did not undergo any significant conformational changes.

**Figure.2.**
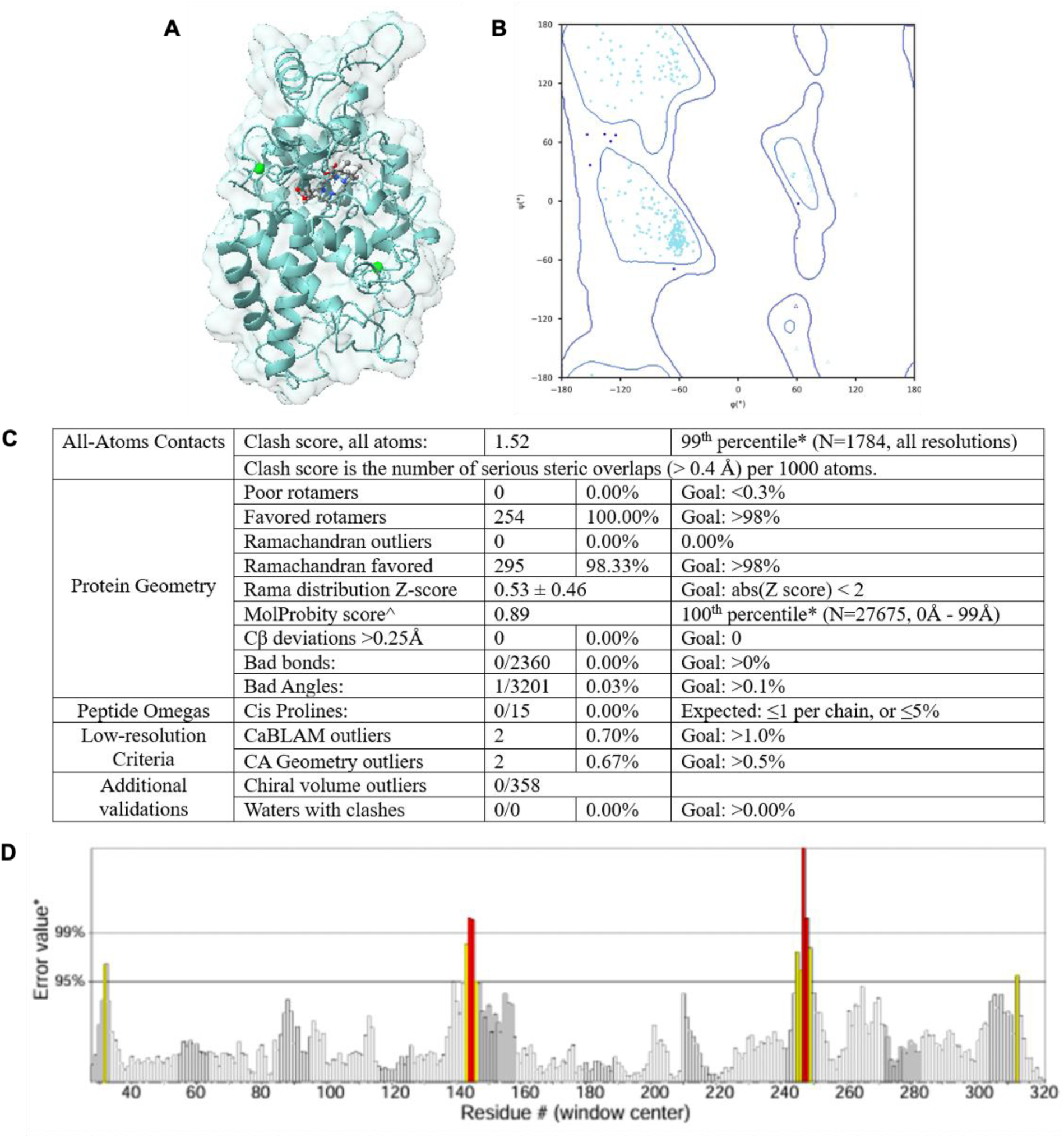
Prediction and validation of TomEP 3D structure. (A) Cartoon structure of TomEP, (B) Ramachandran plot of the predicted 3D structure obtained from MolProbity server, (C) MolPobity results and (D) ERRAT error values graph of TomEP.

**Figure.3.**
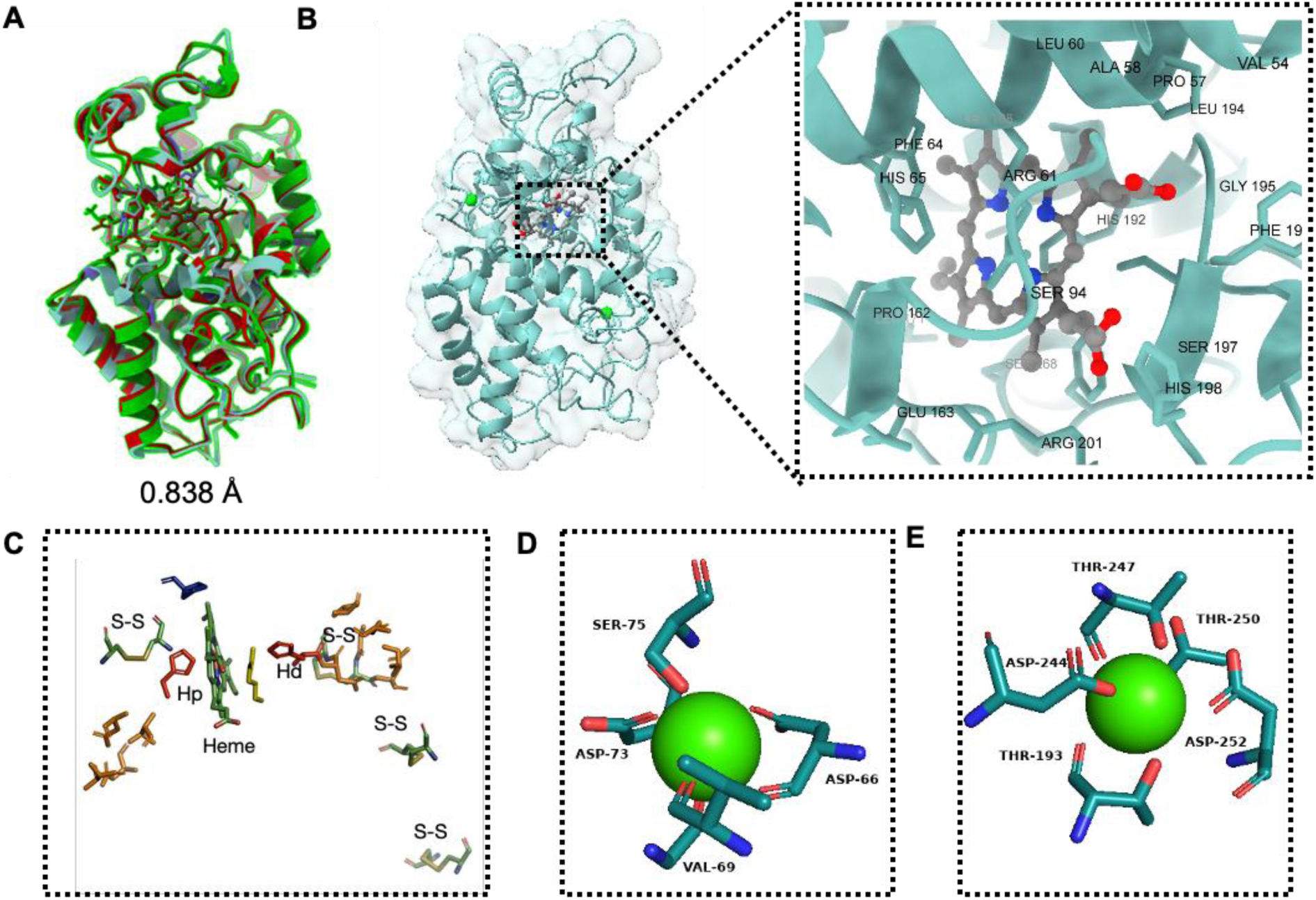
3D structure, active and binding sites of TomEP. The 3D structure of TomEP was predicted using alphfold 3.0 and PyMOL was used to show the distal and proximal domain, heme group, and two calcium-binding sites **(A)** Alignment of TomEP 3D structure with HRP-C (Horseradish peroxidase isoform C) and GvEP1 (Extensin Peroxidase from grape vine) **(B)** TomEP 3D structure **and** heme active site, **(C)** Catalytic center of TomEP **(D)** TomEP distal calcium binding site and **(E)** proximal calcium binding site.

The TomEP 3D structure was further assessed using Ramachandran plot, MolProbity, Errat, ProSA, ProQ and PROCHECK programs. The MolProbity score, which compares the backbone dihedral angles phi and psi of amino acid residues with Ramachandran plot, was 0.89, with 98.33% of the angles matching the feasible locations in the Ramachandran plot. No rotamer outliers were found (0.00%) with no C-beta deviation. There were no bad bonds (0/2538) with only one bad angle (1/3443) found (Fig. 2B and 2C). The ERRAT analysis showed that the TomEP structure was of high quality with an overall quality factor value of 96.26% (Fig. 2D), while the ProQ LG score was 7.29 and ProSA Z-Score was -8.17. The TomEP structure also passed the PROCHECK analysis with 92.4% of amino acids found in the most favored region, 7.2% additionally allowed, 0.4% generously allowed, while there were no amino acids in disallowed regions. Figure 2 and Table S5 summarize the validation analysis of TomEP’s 3D structure. This modeled TomEP 3D structure was used for subsequent molecular docking and MD simulations and compared to HRP-C and *Gv*EP1.

### The active site of TomEP shares core features with other class III peroxidases but has an expanded binding cavity and distinct amino acid environment

All class III peroxidases, including TomEP, have a heme cofactor occupying a central location (Fig. 3B and 3C), with a distal domain (above the location of the heme) and the proximal domain below it form the active site. The Histidine (His-192) located in the proximal domain is conserved in all peroxidases and may play a role in the coordination of the heme. The binding domain has four disulfide bridges, and two calcium ions (distal and proximal Ca) which are also characteristics of class III peroxidases (Fig. 3B and 3C). The distal calcium ion interacts with Asp66, Val69, Asp73 and Ser75 residues through an extensive hydrogen-bonding network (Fig. 3D). In parallel, the proximal calcium ion forms hydrogen bonds with Thr193, Asp244, Thr247, Thr250, and Asp252 (Fig. 3E).

The active site of TomEP was compared with HRP-C and *Gv*EP1 peroxidases using PyMOL and ChimeraX. The cofactor heme group is bound to the protein through a coordinate bond from the heme to the proximal His170 in HRP, His196 in *Gv*EP1 and His192 in TomEP (Fig. 4A). The key catalytic residues His and Arg (His42 and Arg38 of HRP, His65 and Arg61 of TomEP and His70 and Arg65 of *Gv*EP1) which are involved in the formation of compound I (Fig. 5A), are at identical positions in all the three peroxidases (Fig. 4A). The entrance or exit to the active sites is placed between opened alpha coils in the three peroxidases (TomEP, HRP-C and *Gv*EP1), (Fig.4 C, 4F, and 4I, respectively). Sterically, it is the only location on the protein’s surface that is easily accessible to the substrate. Furthermore, the results showed that most amino acid residues around the binding sites of TomEP (Fig. 4B) and *Gv*EP1 (Fig. 4E) are hydrophobic (Phenylalanine, Proline and Leucine) and less bulky in comparison to those in HRP-C (Fig. 4H). Parameters such as pocket volume, area and the total area and circumference of the pocket opening were also compared using CASTp server. The volume (V) and area (A) of the substrate binding pocket of TomEP were 1574 Å^3^ and 1144.6 Å^2^, respectively, which are comparable to *Gv*EP1 (V:1829.8 Å^3^, A:1068.1Å^2^) but are higher than HRP-C (V:1305 Å^3^, A:954.9 Å^2^). Moreover, the total area (TA) and circumference (C) of the opening were also larger for TomEP (TA:158.3 Å^2^, C:88.23Å) and *Gv*EP1 (TA:225.1 Å^2^, C:121.6Å) in comparison with HRP-C (TA:53.6 Å^2^, C:42.3 Å) (Fig. 4C,D,F,G,I,J and Table S6).

**Fig.4:**
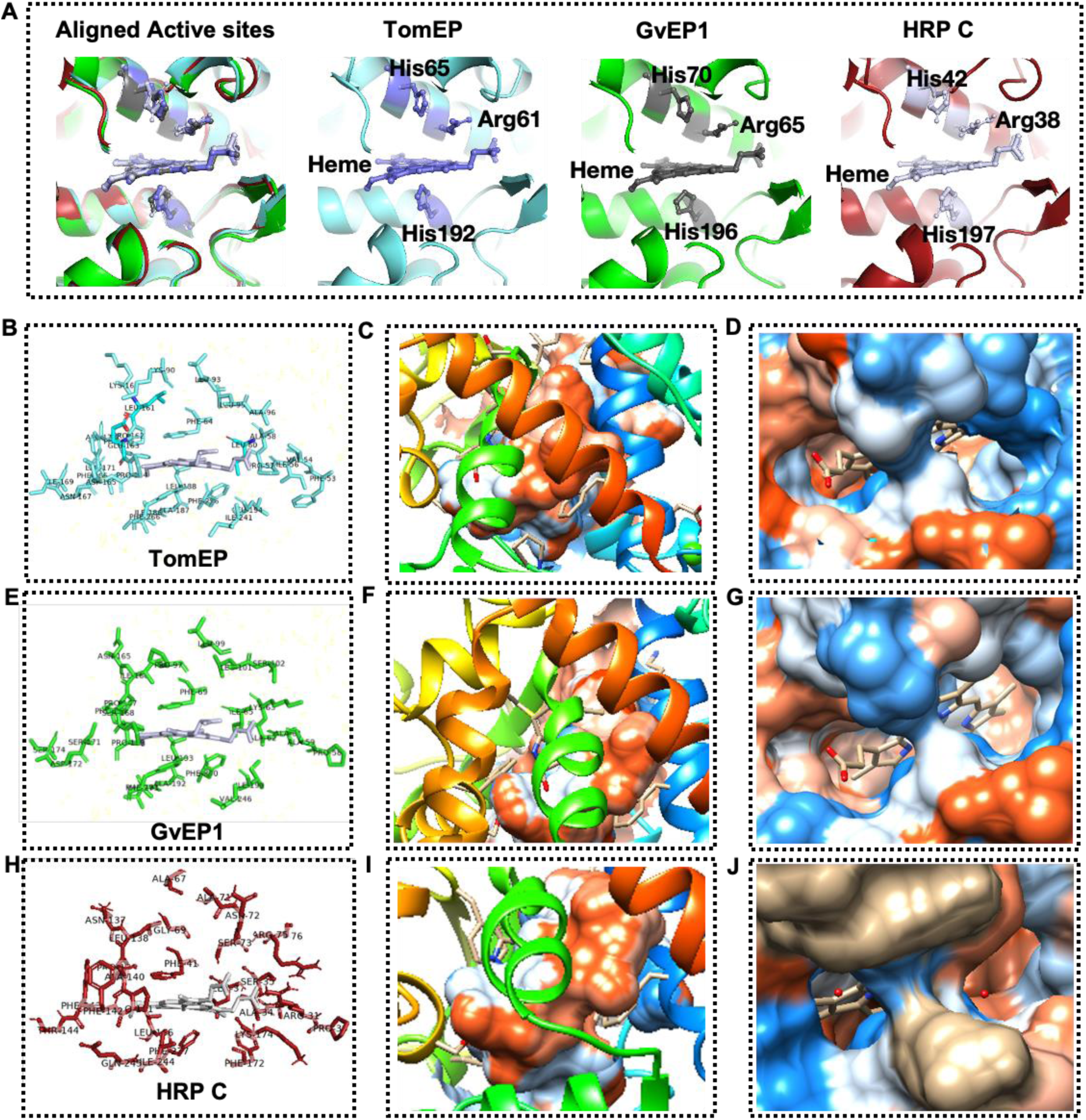
Comparison of Substrate active site and binding pocket of TomEP with other peroxidases. **(A)** Alignment of TomEP with GvEP1 (Extensin Peroxidase from grape vine) and HRP-C (Horseradish peroxidase isoform C) **(B)** Catalytic center of TomEP (Cyan), **(E)** GvEP (Green) and **(H)** HRP-C (Red) protein, (C) TomEP active site cavity, (D) TomEP active site cavity opening, (F) GvEP active site cavity, **(G)** GvEP active site cavity opening, (I) HRP-C active site cavity, (J) HRP-C active site cavity opening.

### The topology of TomEP’s binding sites have similarities to other peroxidases, but are not identical

X-ray crystallographic structures of some peroxidases in complex with small ligands are available in the protein data bank. However, there is no crystal structure of EPs with any large ligand resolved. The binding fitness of [-Y-X-Y-] motifs to the catalytic site of TomEP was evaluated using molecular docking followed by MD simulation. The results were then compared to the available crystal structure of the HRP-C- BENZHYDROXAMIC ACID (2ATJ) [56]. Docking was performed using PyRx software to determine binding conformations, binding residues and binding affinity of the substrate ligands ([-Y-X-Y-] and derivatives (i.e., IDT, Pul and DiIDT) in complex with TomEP. The complexes having the highest negative binding values (i.e., high affinity) were selected for further analysis and identification of amino acids that interact with the ligands. These results were compared to experimentally observed HRP-C- BENZHYDROXAMIC ACID (2ATJ) crystal structure) to highlight differences in the ligand binding pocket between the two enzymes [60]. The docked complexes of TomEP+[-Y-X-Y-] were compatible with the binding of benzhydroxamic acid to HRP-C (Fig. 5A and 5B). In both structures, the substrates were oriented in accordance with previously presented results of the active site accessibility (Fig. 4C). Despite the high similarity, the topology of the two binding sites is not identical, which could be explained by the difference in the substrate structures and the size of the binding sites. The position of benzhydroxamic acid in complex with HRP-C is close to the heme (Fig. 5A), while in the TomEP-[-Y-X-Y-] complexes (Fig. 5B), the substrate location is next to heme.

**Figure.5.**
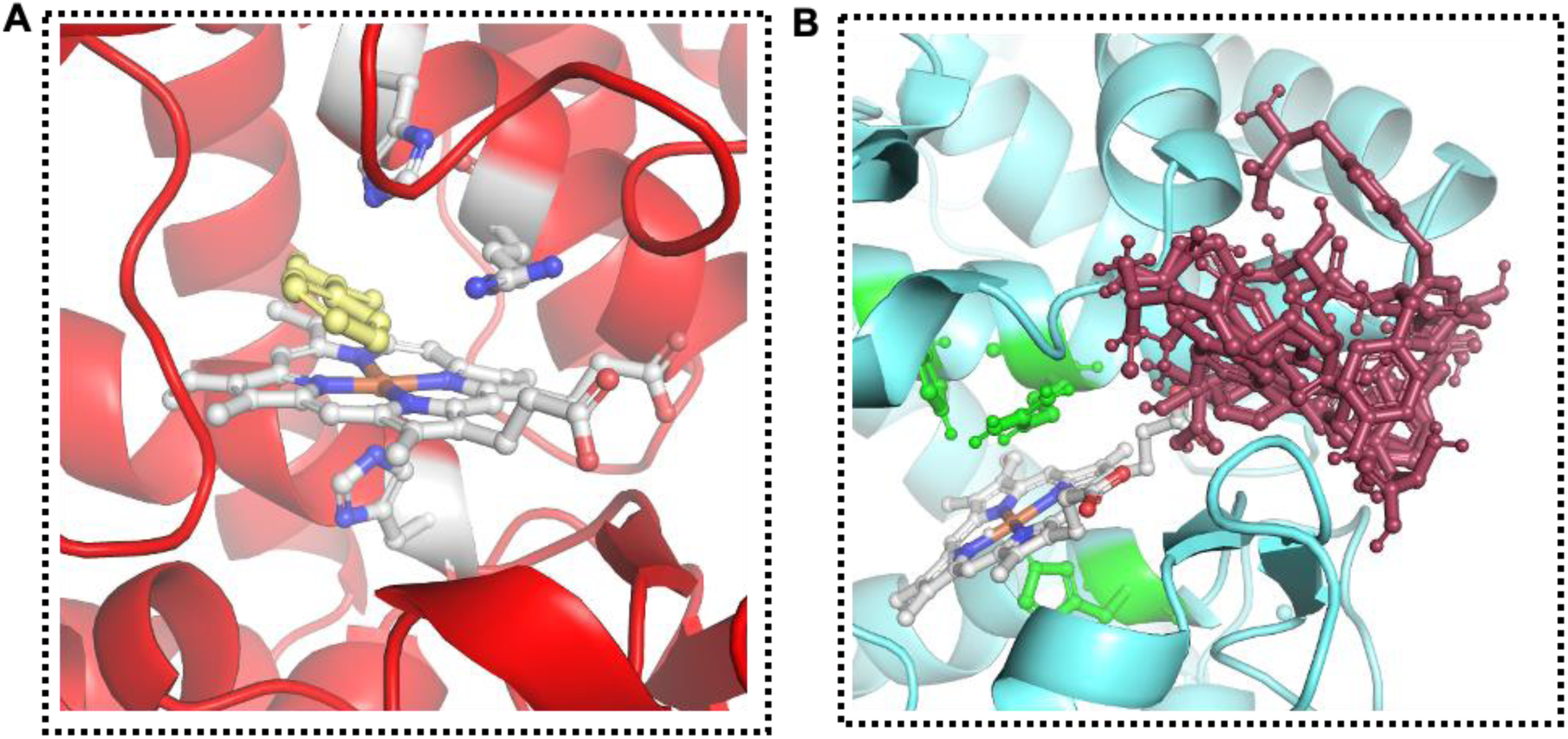
Substrate binding into the catalytic site of HRP-C and TomEP. **(A)** Crystal structure of HRP-BENZHYDROXAMIC ACID (PDB ID: 2ATJ) and **(B)** TomEP docked complexes with -Y-X-Y EXTs ligands and its derivatives.

All of the tested substrate ligands ([-Y-V-Y-], [-Y-K-Y-], [-Y-Y-Y-], Pul, IDT, and Di-IDT) fit nicely inside the binding pocket of TomEP. The binding affinity for [-Y-V-Y-] was -6.6 kcal/mole and it interacted with Ser94, Thr238, Ala240 (through conventional H-bonding), Phe196 (through Pi-Pi stacking), Val54, Ala96, and Ile239 (through Pi-Alkyl) interactions (Fig.6A). For [-Y-Y-Y-], the binding affinity was -6.1 kcal/mole, and interactions occurred *via* Thr52, Ser94, Asp98, Thr238, Ala240, (through conventional H-bonding), Phe196 (through Pi-Pi stacking), Val96 (through Pi-Alkyl), Ala96 (through Pi-Sigma) and His198 (through carbon H-bonding) (Fig. 6B). The binding affinity for Pulcherosine (Pul) was -6.5 kcal/mole and it interacted with Thr52, Phe53, Val54, Leu93, Ser94, Phe196, Arg249 (through conventional H-bonding), and His198 (through carbon H-bonding) (Fig.6C). Results of [-Y-K-Y-] showed a binding affinity of -5.2 kcal/mole, and the interacting residues were Asp98, Thr238, Ser94 (through conventional H-bonding), Phe196 (through Pi-Pi stacking), Ala96 (through Pi-Sigma), and Ala240, Val54, (through Pi-Alkyl) interactions (Fig. S4A). The binding affinity for IDT was -5.9 kcal/mole and the interactions were with Thr52, Leu93, Ser94, Gly97, Ala240 Arg249 (through conventional H-bonding), Phe196 (through Pi-Pi stacking) and Ala96 (through Pi-Alkyl) (Fig.S4B). Finally, for Di-IDT, the binding affinity was -6.1 kcal/mole, and the interactions were with Thr52, Phe53, Val54, Leu93, Gly97, Asp98, Ala240 (through conventional H-bonding), Phe196 (through Pi-Pi stacking), His198 (through Pi Cation) and Ala96 (through carbon hydrogen bonding) interactions (Fig. S4C). These results revealed that a set of key amino acids in TomEP interact with substrate ligands and may contribute to the specificity of substrate recognition and binding. Notably, a few of these amino acids (i.e, Val54, Ser94, Ala96 and Phe196) are conserved across all the docked complexes suggesting their potential role in TomEP catalytic mechanism. Together, these results provide an insight into the interaction of TomEP with [-Y-X-Y-] substrate ligands and provide a frame for future functional and mutational studies.

**Fig.6:**
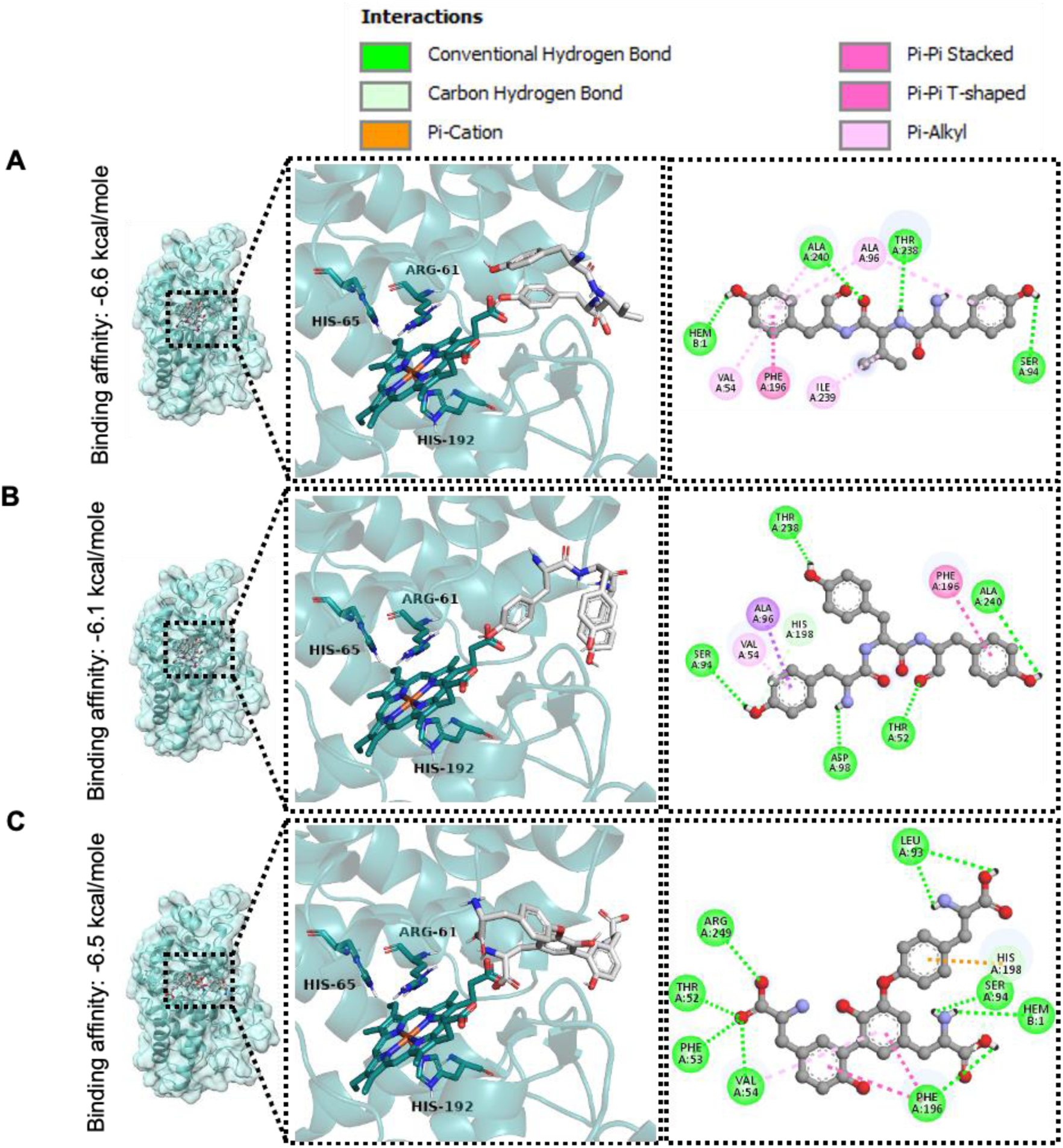
Docking pose and binding affinity of docked EXTs ligands with TomEP. Docked complexes of (A) YVY, (B) YYY and (C) Pulcherosine in TomEP. The left image depicts 3D structure of TomEP with bound ligand while the right image depicts the ligand bound in active site and 2D interactive plot of ligands contacts.

### Molecular dynamics simulations demonstrate formation of stable TomEP+[-Y-X-Y-] complexes

The stability of molecular docking results was evaluated using MDS. Molecular docking showed highly favorable binding interactions between TomEP and the studied substrate ligands. However, it should be noted that the stability of the protein-ligand interaction relies upon the molecular interactions and the surrounding solvent conditions. Molecular docking is computationally fast, but it does not account for the solvent and various entropic effects. In certain cases, substrate ligands that exhibit high molecular interactions and docking scores may not bind effectively to the protein in an experimental setup [57, 58]. To validate the docking results, we performed fully atomistic MDS using Desmond package of Maestro (Schrodinger). The main goal of these simulations was to determine if the substrate ligands would release from the active site during the simulation time, or if the protein undergoes a conformational change due to the bound ligand. The simulation confirmed the stability of the binding, as all the substrate ligands ([-Y-V-Y-], [-Y-K-Y-], [-Y-Y-Y-], Pul, IDT, and Di-IDT) remained stable in complexes with heme during the 100 ns (Fig. 7). The substrates’ binding enhanced TomEP’s stability, which became even more stable in comparison with the simulation in the absence of substrates (Fig. 7B, S2C and S2D). We traced Cα-RMSD values over time for 100 ns (Fig. 7A) and observed that the RMSD trajectory of TomEP swiftly reached a stable equilibrium within 1 ns of simulation for all the protein-substrate ligands complexes. The complexes maintained an average RMSD change of 0.10±0.02 Å for [-Y-Y-Y-] and [-Y-V-Y-], 0.01±0.02Å for [-Y-K-Y-], 0.02±0.02Å for Pul and DiIDT, while IDT showed 0.03±0.02Å change over the 1-100 ns time frame (Fig. 7A). The RMSF results also revealed that the secondary structures of the protein remain stable throughout the 100 ns simulation for all the complexes. In fact, a significant reduction in RMSF values was noticed, showing that the binding of substrate reduces the fluctuation, and the protein structure becomes more stable after substrate binding (Fig. S2D). The average RMSF values observed for ligand over TomEP complexes were 0.078±0.02Å ([-Y-V-Y-]), 0.077±0.02Å ([-Y-Y-Y-]), 0.073±0.02Å ([-Y-K-Y-]), 0.076±0.02Å (IDT), 0.095±0.02 (Pul), and 0.088±0.02Å (DiIDT) (Fig. 7B). Taken together, these results support the conclusion that the docked complexes (TomEP+[-Y-X-Y-] and derivatives) were stable, which should allow the catalysis to take place with minimal structural deviation and fluctuation and support the strength of predicted 3D structure of TomEP. In a nutshell, MDS results validate the binding fitness, and provide dynamic insights on conformational stability that underpin TomEP function.

**Fig.7:**
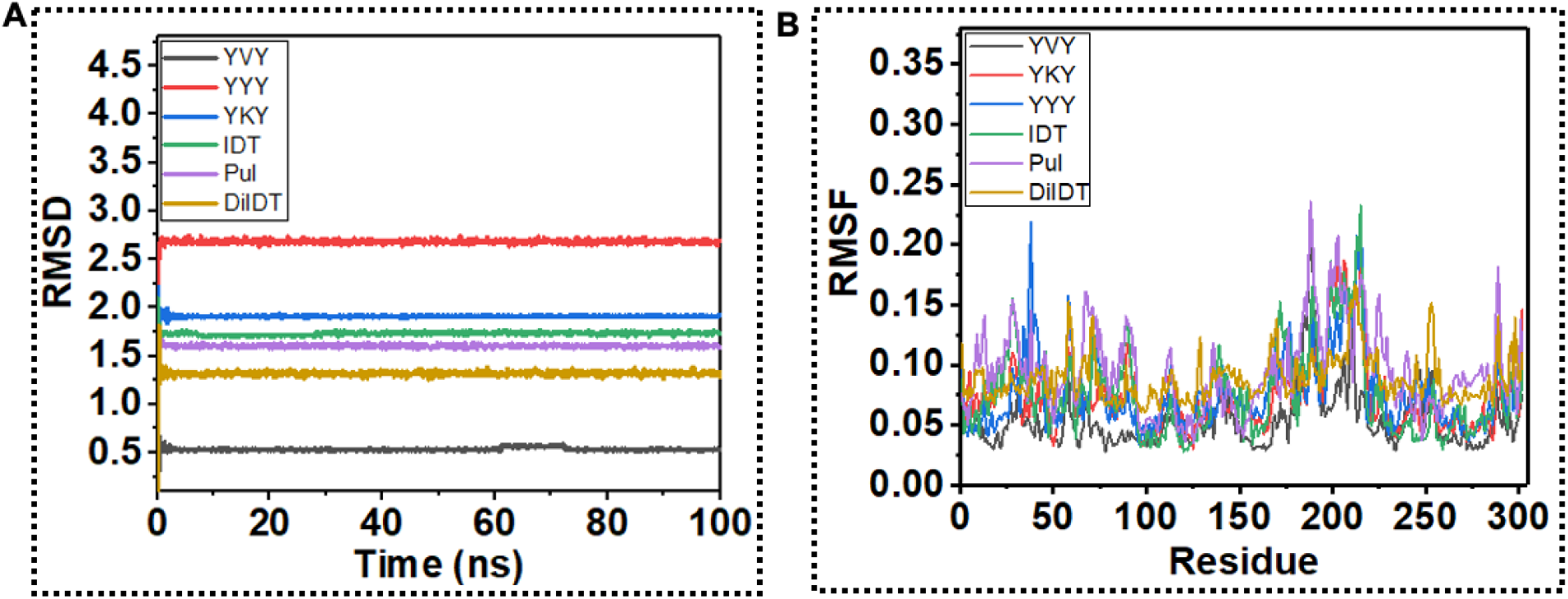
Molecular dynamics simulation results of TomEP and EXTs ligands complexes. TomEP EXT ligand complexes’ (A) Root mean square deviations (RMSD) and (B) Root mean square fluctuation during molecular dynamics simulations. Colors represent different protein ligand complexes.

## Discussion

Although EXTs comprise only about 10% of the primary cell wall [5] their crosslinking is required for proper cell wall assembly, particularly during cell division, embryo formation and seeds germination [11, 59, 60]. The tripeptide motif [-Y-X-Y-] of EXT monomers can become covalently crosslinked to each other forming, IDT, Pul and di-IDT [61, 62]. EPs are enzymes that catalyze these crosslinking reactions. Several peroxidase candidates have been described in various plant species that are associated specifically with crosslinking of EXTs in *in vitro* studies (LEP1 (Lupin EP), *Gv*EP1, and FBP1 (French bean peroxidase)) or immunolocalization (PRX09 and PRX40) [25–27, 63]. However, their biochemical mechanisms remain unexplored because of inadequate information on the active site and key amino acid residues that interact with the prospective substrate ligands. Also, what makes certain peroxidases (EPs) more specific towards EXTs is still unclear. To address these limitations, a predictive *in silico* approach along with structural analysis can help determine the characteristics of the active sites of these enzymes, including substrate binding fitness and possible catalytic mechanism. Understanding the structural basis of the biochemical mechanism would allow engineering EPs for crops improvement and/or generating novel materials using EXTs.

In the absence of crystal structure of EPs, computational approaches such as prediction of 3D protein structures, molecular docking, and molecular dynamics simulations are an excellent alternative to advance our understanding of the structural basis of enzyme catalytic activity [64]. These computational approaches have also been widely used in studying other peroxidases [32, 33, 34, 35, 65–67]. However, no extensive computational studies of EPs have been conducted. Several studies have shown TomEP to be a potential candidate to catalyze crosslinking of [-Y-X-Y-] linkages, but the molecular mechanism is not established yet [23, 24, 30]. This study fills this gap by investigating the binding interactions and affinity of [-Y-X-Y-] substrate ligands and its derivatives within the active site of TomEP. The prediction and assessment of 3D structures are pre-requisites for molecular docking and molecular dynamics simulations. In this study, we predicted the secondary and 3D structures of TomEP, its physicochemical properties, catalytic cavity and protein-substrate ligands interactions. The stability of docked substrate ligands was further confirmed through isothermal isobaric ensemble molecular dynamics simulations for 100 ns [43]. The predicted structure of TomEP contains the key characteristics of class III peroxidases, such as a domain that contains heme, distal and proximal His, and the presence of eight cysteine residues involved in four disulphide bridges [68]. In a first step, the predicted 3D structure of TomEP was evaluated and confirmed with various bioinformatics tools such as PROCHECK ERRAT, ProSA, ProQ and Ramachandran plot. The results confirmed that our predicted 3D structure was of high quality and structurally reliable. Next, we compared the 3D structure of TomEP with publicly available structures of known peroxidases by searching DALI database. The search showed that all the closest structures were also class III peroxidases, giving further validation of the predicted 3D structure of TomEP. Furthermore, MDS was conducted for 100 ns to assess the structural stability of TomEP [69]. The molecular dynamics simulation results showed that the TomEP predicted 3D structure was stable for 100 ns.

Having a reliable 3D structure of TomEP allowed us to conduct detailed analyses of the substrate binding pocket and the fitness of the binding of the substrates and derivatives. The rational is that the substrate binding pocket of an EP must be large enough to accommodate EXT substrate. To verify this hypothesis, the binding pocket of TomEP was analyzed and compared with other characterized peroxidases such as *Gv*EP1 (a known EP) and a peroxidase that has a low affinity for EXT substrates such as HRP-C, using PyMOL, ChimeraX, CASTp programs. Using CASTp program, (which measures the area and volume of binding pocket) [70], and PyMOL and ChimeraX, we showed that all the three peroxidases (TomEP, *Gv*EP and HRP-C) have similar characteristics in terms of overall protein fold and site structure. This structural similarity is important because of their highly similar mechanism of functioning [31, 71]. However, our analyses showed that TomEP and *Gv*EP1 have more hydrophobic residues compared to HRP-C (low affinity for EXT substrates). Also, the volume and area of the binding pocket and opening of TomEP and *Gv*EP1 were significantly higher compared to HRP-C. Features such as enlarged binding pockets and larger size of the entrance could reflect the unique structural and functional adaptation of EPs for longer peptide chain with [-Y-X-Y-] motifs, which make them more accessible to the active site. In addition, the docking and MDS also support this hypothesis by demonstrating that the substrate ligands are stable inside the binding cavity.

Our analysis of the binding pocket revealed Val54, Ser94, Ala96 and Phe196 residues as key interacting amino acids with numerous types of interactions. The catalytic reaction of EPs can go through either of two complementary cycles: the peroxidative cycle (Fig. S5A) or the hydroxylic cycle (Fig. S5B). The peroxidative cycle is responsible for crosslinking of EXT monomers [26, 72]. The mechanism of the peroxidative cycle consists of the oxidization of the native ferric resting form by H_2_O_2_ and the formation of an unstable intermediate called compound I (CompI, Fig. S5A), which oxidizes the electron donor substrate (IDT^EXT1^) to become CompII releasing a free radical. CompII is further reduced by a second substrate (IDT^EXT2^) regenerating the Fe III state of the enzyme and releasing another free radical. The two oxidized forms of IDT (EXT1 and EXT2) produce DiIDT (Mishler-Elmore, 2020) (Fig. S5A). Among the above-mentioned residues, Ser94 is more nucleophilic having a hydroxyl (-OH) group on its side chain and might play a potential role in electron transfer during peroxidative cycle.

Although this study is based purely on *in silico* analysis, it provides insights about the structural basis the specificity of TomEP and GvEP1 towards EXTs. However, several limitations must be acknowledged. The predicted 3D structure of TomEP (alone and in complex with ligands) should be validated experimentally by resolving the structure through crystallography. Furthermore, experimental validation of Val54, Ser94, Ala96 and Phe196 residues’ involvement in catalysis is necessary and can be achieved through point mutagenesis and *in vitro* assays. This work can be used as a working frame for future *in vivo* experimental design.

## Conclusion

The current study provides structural and functional features of TomEP through molecular modeling analyses. The 3D structure of TomEP perfectly superimposed to the structures of known peroxidases from class III. A total of 6 different [-Y-X-Y-] substrate ligands have been assessed to evaluate the binding fitness with TomEP. All the substrate ligands tested bound stably inside the TomEP binding pocket. Our results provide the structural basis that strongly support TomEP as a suitable candidate for crosslinking of [-Y-X-Y-] substrate ligands and derivatives. These structural features were not present in other class III peroxidases.

## Abbreviation

EPs: Extensin peroxidases
EXT: Extensin
3D: Three-dimensional
TomEP: Tomato EP
-Y-X-Y-: Tyrosine-X-Tyrosine
-YKY-: Tyrosine-Lysine-Tyrosine
-YVY-: Tyrosine-Valine-Tyrosine
-YYY-: Tyrosine-Tyrosine-Tyrosine
IDT: Isodityrosine
Pul: Pulcherosine
diIDT: Di-Isodytirosine
Gal: galactosyl
Ara*f*: Arabinofuranosyl
AGPs: Arabinogalactan proteins
Hyp: Hydroxyproline
MDS: Molecular Dynamics Simulation
pI: Isoelectric point
EC: Extinction Coefficient
II: Instability index
SOPMA: Self-Optimized Prediction Method with Alignment
SAS: Sequence Annotated by Structure
GRAVY: Grand average of hydropathicity
HRP-C: Horse Radish Peroxidase
*Gv*EP1: Grapevine extensin peroxidase
OPLS3: Optimized Potentials for Liquid Simulations
RMSD: Root Mean Squared Deviation
RMSF: Root Mean Square Fluctuations

## Declarations

**Clinical Trial Number:** Not applicable

**Ethics declarations:** Not applicable

**Ethics approval and consent to participate:** Not applicable

**Consent for publication:** Not applicable

## Availability of data and materials

All data generated in this study (including files for 3D alpha fold structures, ligand structures, dock complexes, and the molecular dynamics simulations) are available upon request from the authors.

## Contributions

MH, SS, and AF conceptualized the study. MH, SS, AF and ZH helped design the experimental work. ZH performed and collected the experimental work with input from SS. Data analysis was performed by MH, SS, AF, and ZH. ZH prepared the data, the supplementary materials, and manuscript. MH, AF, and SS assisted with manuscript writing/editing and overall project supervision.

## Acknowledgements

We would like to thank Miss Shehnaz, for contributions to manuscript preparation. We are also thankful to Allan Regunton, Abdul Hakeem Al Bulushi, Maryam Muzaffar, Fitrat Ullah, Samia Nawaz and Mohsin Ali Nasir for their feedback and suggestions.

