## supplemental File 1l for "*In silico* analysis reveals the structural basis of TomEP specificity, a tomato extensin peroxidase"

### Slide 1
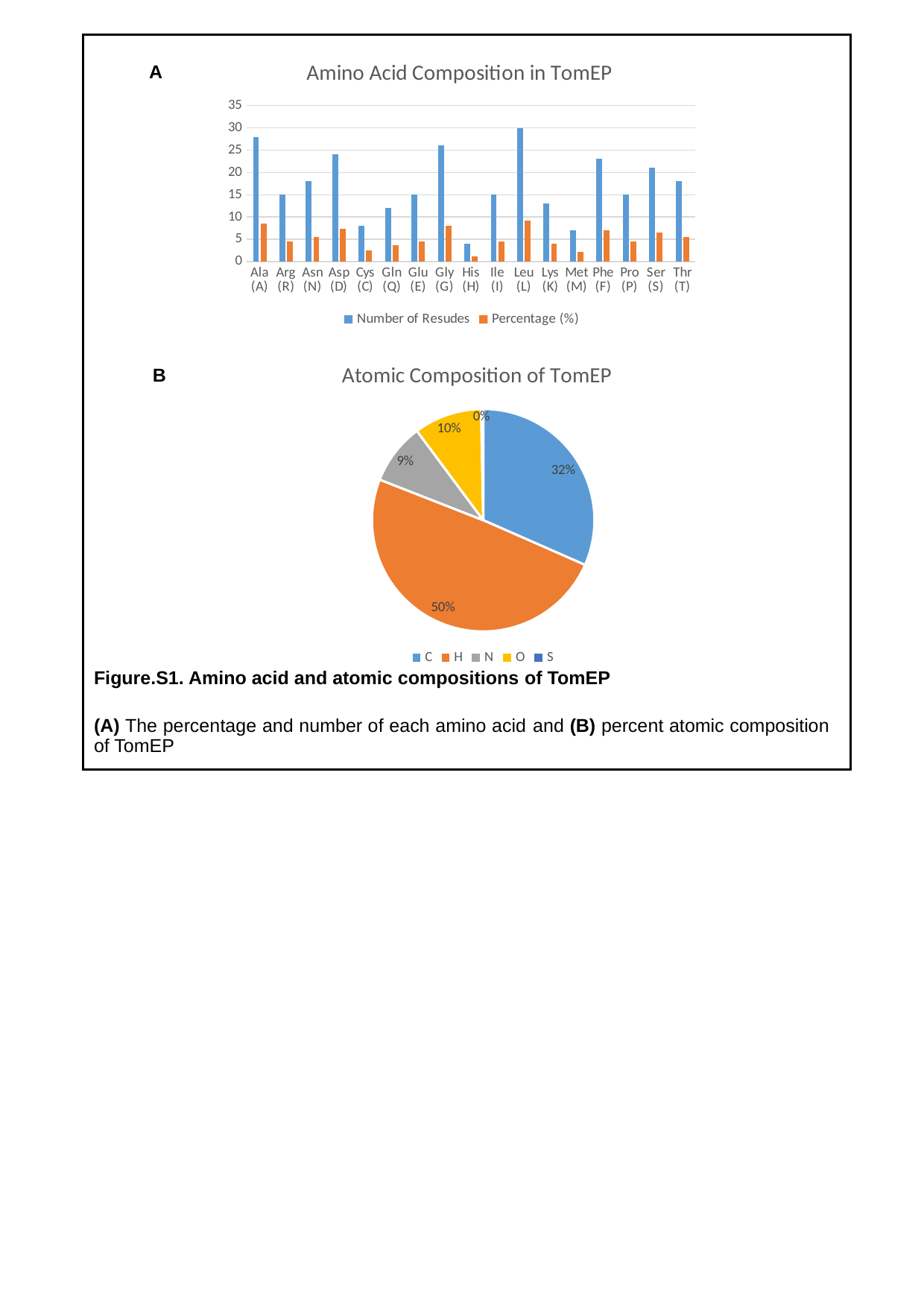

#### Chart: Amino Acid Composition in TomEP
| Category | Number of Resudes | Percentage (%) |
|---|---|---|
| Ala (A) | 28.0 | 8.6 |
| Arg (R) | 15.0 | 4.6 |
| Asn (N) | 18.0 | 5.5 |
| Asp (D) | 24.0 | 7.4 |
| Cys (C) | 8.0 | 2.5 |
| Gln (Q) | 12.0 | 3.7 |
| Glu (E) | 15.0 | 4.6 |
| Gly (G) | 26.0 | 8.0 |
| His (H) | 4.0 | 1.2 |
| Ile (I) | 15.0 | 4.6 |
| Leu (L) | 30.0 | 9.2 |
| Lys (K) | 13.0 | 4.0 |
| Met (M) | 7.0 | 2.2 |
| Phe (F) | 23.0 | 7.1 |
| Pro (P) | 15.0 | 4.6 |
| Ser (S) | 21.0 | 6.5 |
| Thr (T) | 18.0 | 5.5 |A
#### Chart: Atomic Composition of TomEP
| Category | Percentage (%) |
|---|---|
| C | 0.32 |
| H | 0.5 |
| N | 0.09 |
| O | 0.1 |
| S | 0.003 |B
Figure.S1. Amino acid and atomic compositions of TomEP
(A) The percentage and number of each amino acid and (B) percent atomic composition of TomEP

### Slide 2
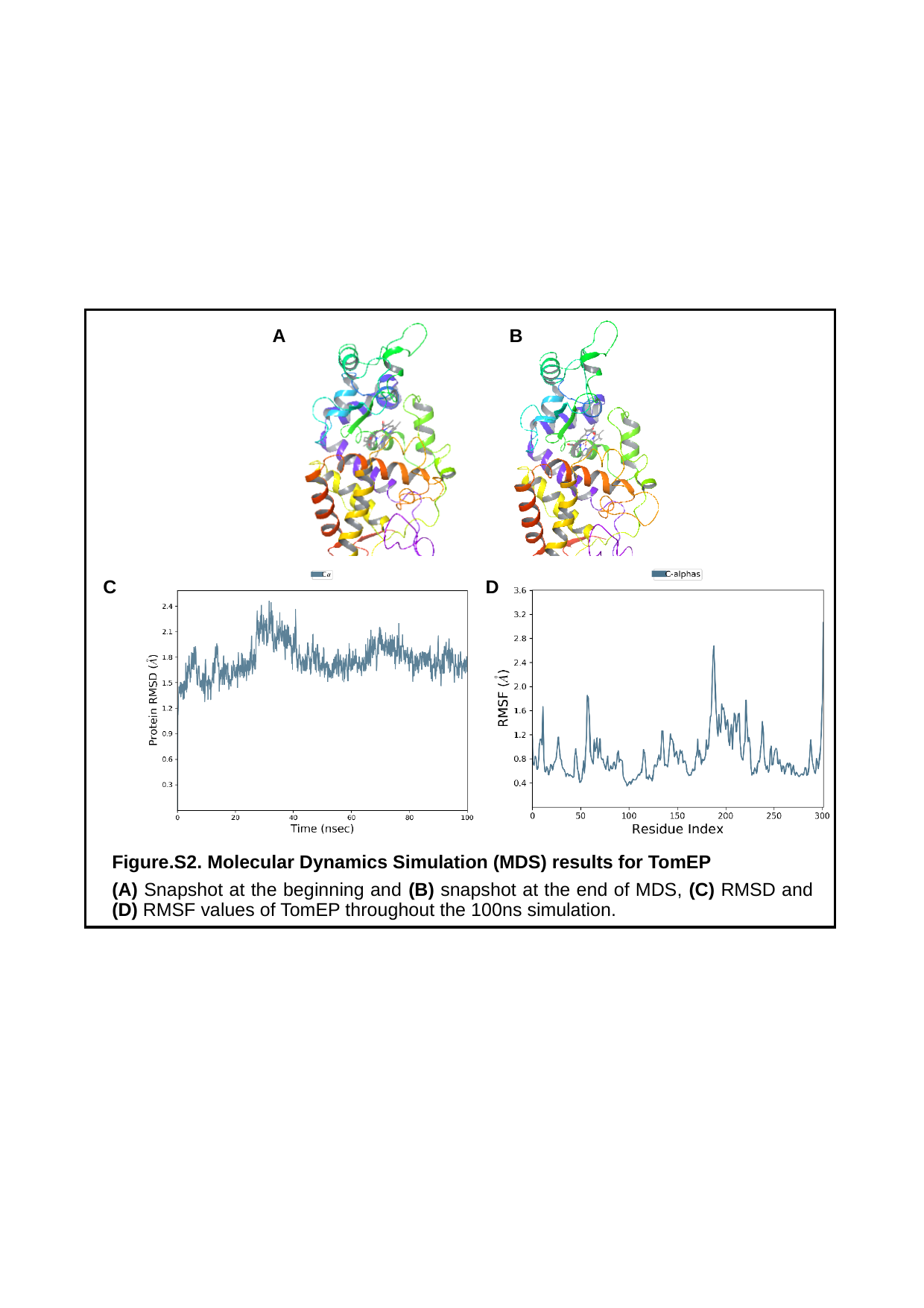

B
A
C
D
Figure.S2. Molecular Dynamics Simulation (MDS) results for TomEP
(A) Snapshot at the beginning and (B) snapshot at the end of MDS, (C) RMSD and (D) RMSF values of TomEP throughout the 100ns simulation.

### Slide 3
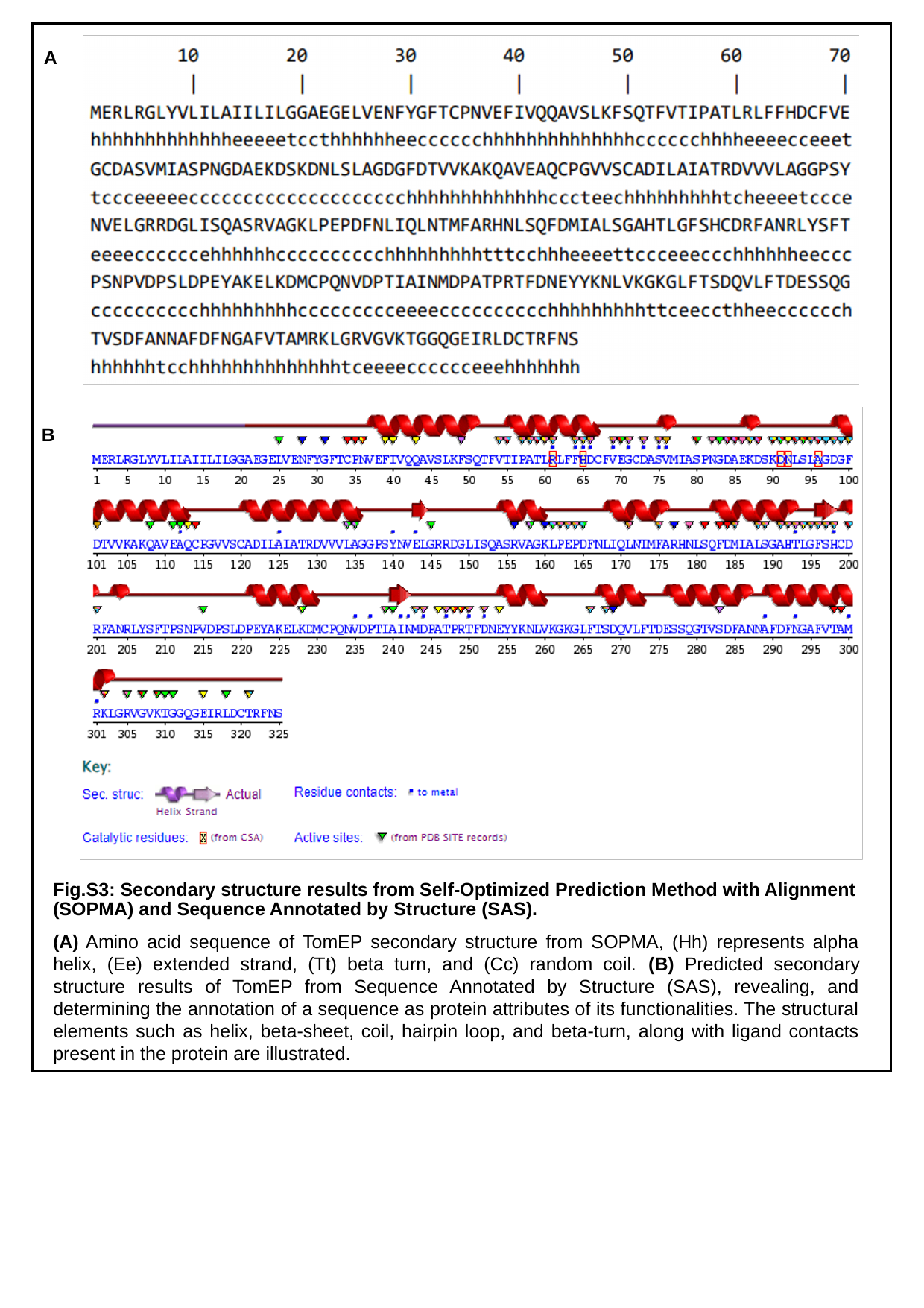

A
B
Fig.S3: Secondary structure results from Self-Optimized Prediction Method with Alignment (SOPMA) and Sequence Annotated by Structure (SAS).
(A) Amino acid sequence of TomEP secondary structure from SOPMA, (Hh) represents alpha helix, (Ee) extended strand, (Tt) beta turn, and (Cc) random coil. (B) Predicted secondary structure results of TomEP from Sequence Annotated by Structure (SAS), revealing, and determining the annotation of a sequence as protein attributes of its functionalities. The structural elements such as helix, beta-sheet, coil, hairpin loop, and beta-turn, along with ligand contacts present in the protein are illustrated.

### Slide 4
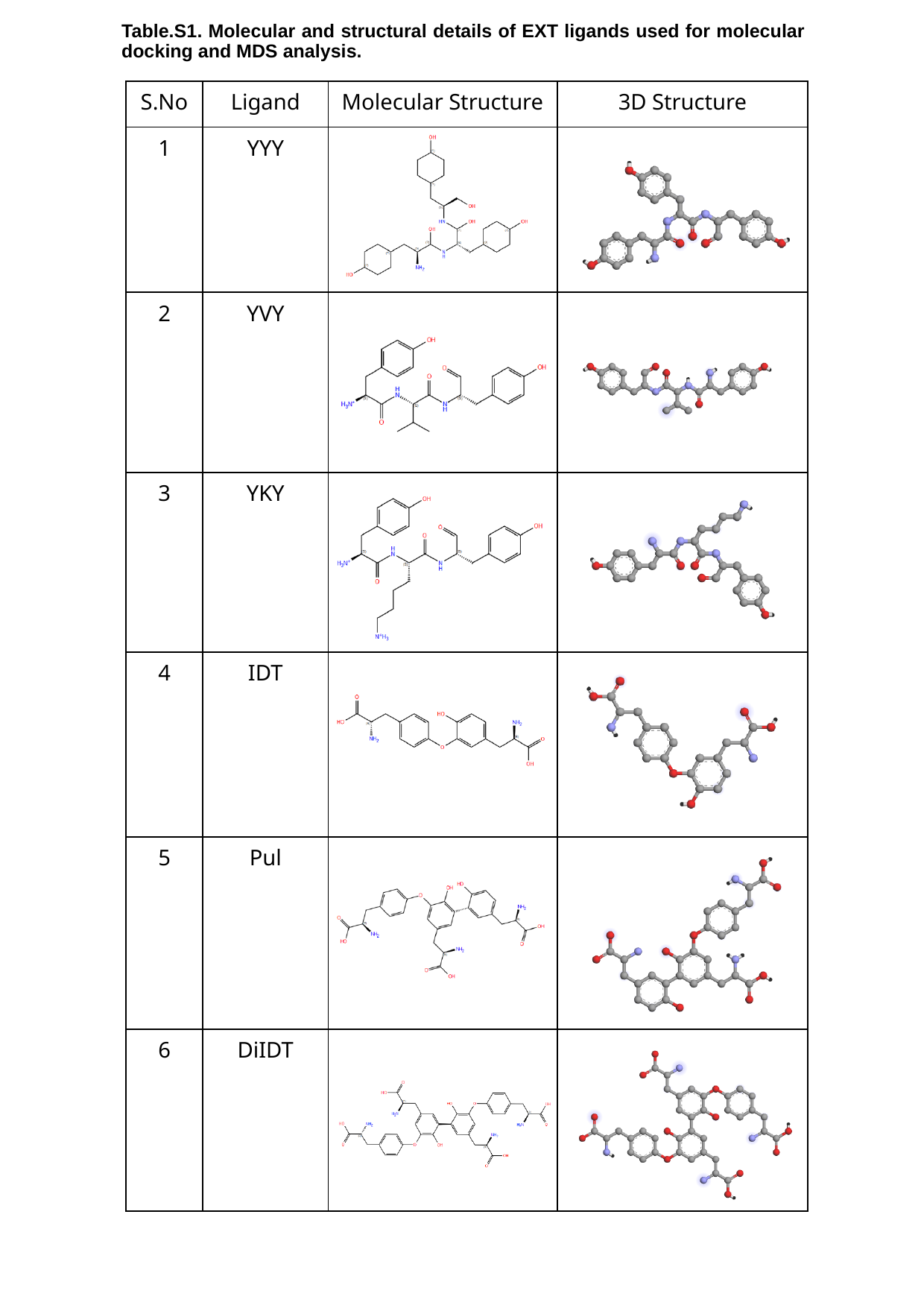

Table.S1. Molecular and structural details of EXT ligands used for molecular docking and MDS analysis.
| S.No | Ligand | Molecular Structure | 3D Structure |
| --- | --- | --- | --- |
| 1 | YYY | | |
| 2 | YVY | | |
| 3 | YKY | | |
| 4 | IDT | | |
| 5 | Pul | | |
| 6 | DiIDT | | |

### Slide 5
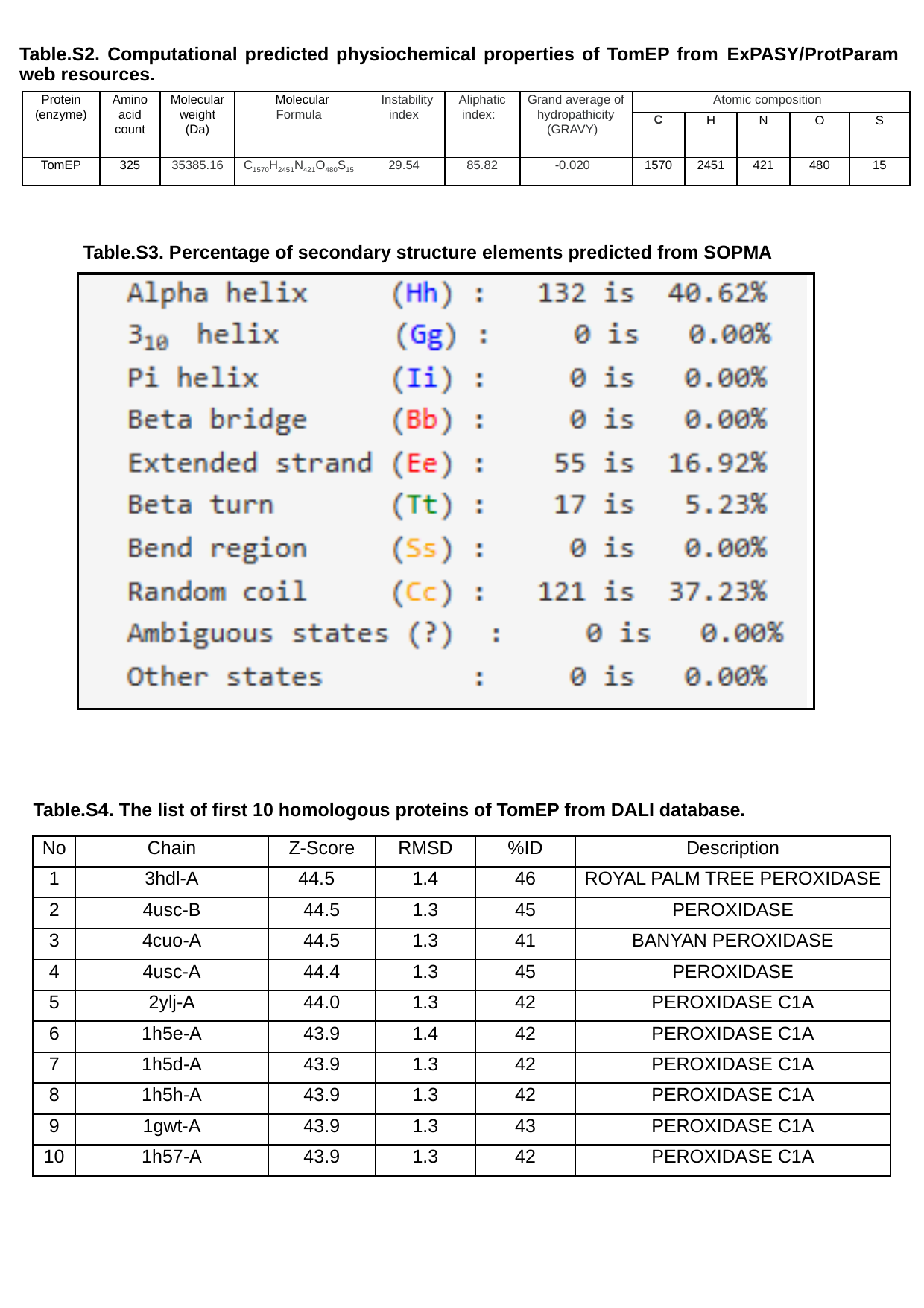

Table.S2. Computational predicted physiochemical properties of TomEP from ExPASY/ProtParam web resources.
| Protein (enzyme) | Amino acid count | Molecular weight (Da) | Molecular Formula | Instability index | Aliphatic index: | Grand average of hydropathicity (GRAVY) | Atomic composition | | | | |
| --- | --- | --- | --- | --- | --- | --- | --- | --- | --- | --- | --- |
| | | | | | | | C | H | N | O | S |
| TomEP | 325 | 35385.16 | C_1570_H_2451_N_421_O_480_S_15_ | 29.54 | 85.82 | -0.020 | 1570 | 2451 | 421 | 480 | 15 |
Table.S3. Percentage of secondary structure elements predicted from SOPMA
Table.S4. The list of first 10 homologous proteins of TomEP from DALI database.
| No | Chain | Z-Score | RMSD | %ID | Description |
| --- | --- | --- | --- | --- | --- |
| 1 | 3hdl-A | 44.5 | 1.4 | 46 | ROYAL PALM TREE PEROXIDASE |
| 2 | 4usc-B | 44.5 | 1.3 | 45 | PEROXIDASE |
| 3 | 4cuo-A | 44.5 | 1.3 | 41 | BANYAN PEROXIDASE |
| 4 | 4usc-A | 44.4 | 1.3 | 45 | PEROXIDASE |
| 5 | 2ylj-A | 44.0 | 1.3 | 42 | PEROXIDASE C1A |
| 6 | 1h5e-A | 43.9 | 1.4 | 42 | PEROXIDASE C1A |
| 7 | 1h5d-A | 43.9 | 1.3 | 42 | PEROXIDASE C1A |
| 8 | 1h5h-A | 43.9 | 1.3 | 42 | PEROXIDASE C1A |
| 9 | 1gwt-A | 43.9 | 1.3 | 43 | PEROXIDASE C1A |
| 10 | 1h57-A | 43.9 | 1.3 | 42 | PEROXIDASE C1A |

### Slide 6
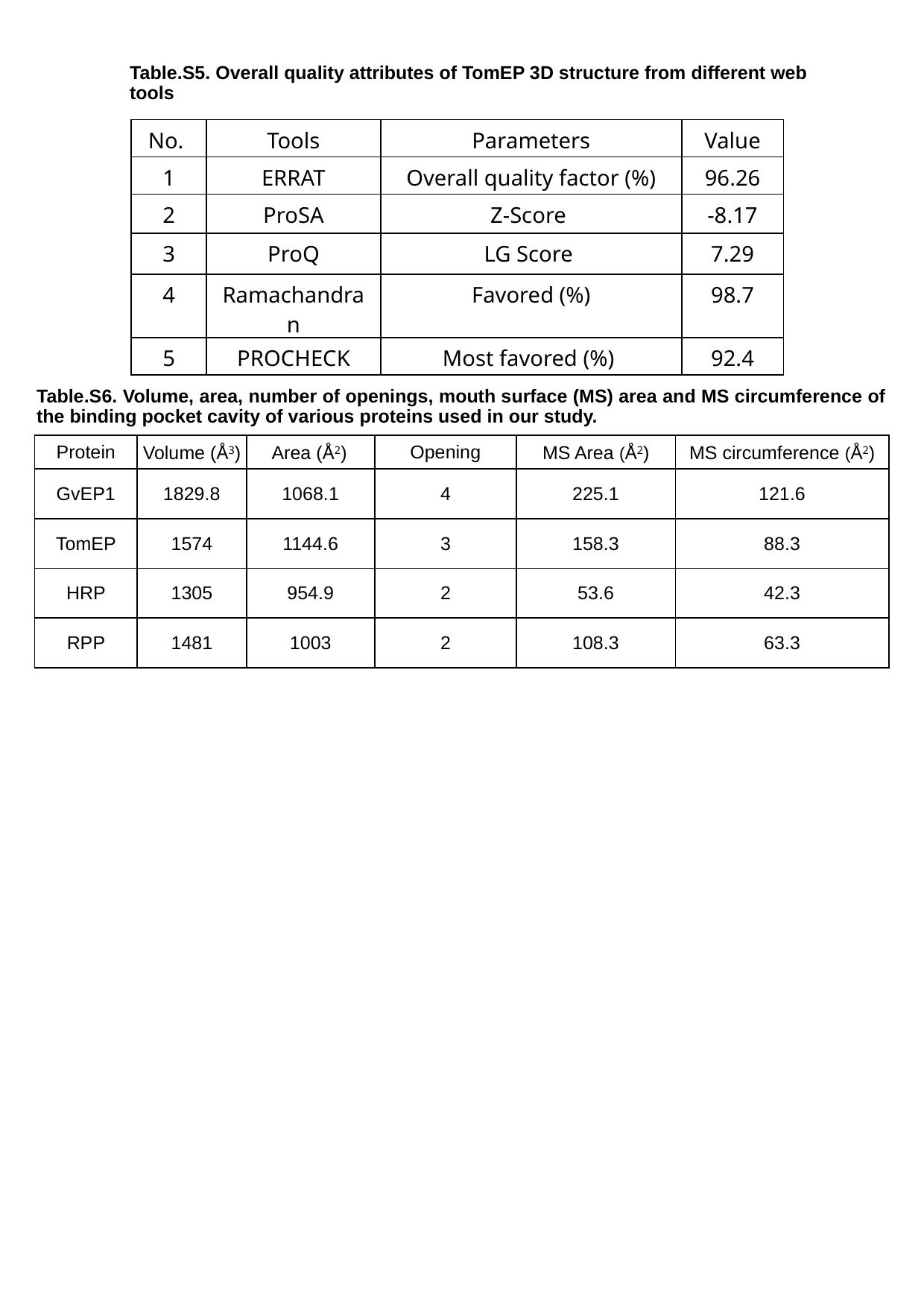

Table.S5. Overall quality attributes of TomEP 3D structure from different web
tools
| No. | Tools | Parameters | Value |
| --- | --- | --- | --- |
| 1 | ERRAT | Overall quality factor (%) | 96.26 |
| 2 | ProSA | Z-Score | -8.17 |
| 3 | ProQ | LG Score | 7.29 |
| 4 | Ramachandran | Favored (%) | 98.7 |
| 5 | PROCHECK | Most favored (%) | 92.4 |
Table.S6. Volume, area, number of openings, mouth surface (MS) area and MS circumference of the binding pocket cavity of various proteins used in our study.
| Protein | Volume (Å^3^) | Area (Å^2^) | Opening | MS Area (Å^2^) | MS circumference (Å^2^) |
| --- | --- | --- | --- | --- | --- |
| GvEP1 | 1829.8 | 1068.1 | 4 | 225.1 | 121.6 |
| TomEP | 1574 | 1144.6 | 3 | 158.3 | 88.3 |
| HRP | 1305 | 954.9 | 2 | 53.6 | 42.3 |
| RPP | 1481 | 1003 | 2 | 108.3 | 63.3 |

### Slide 7
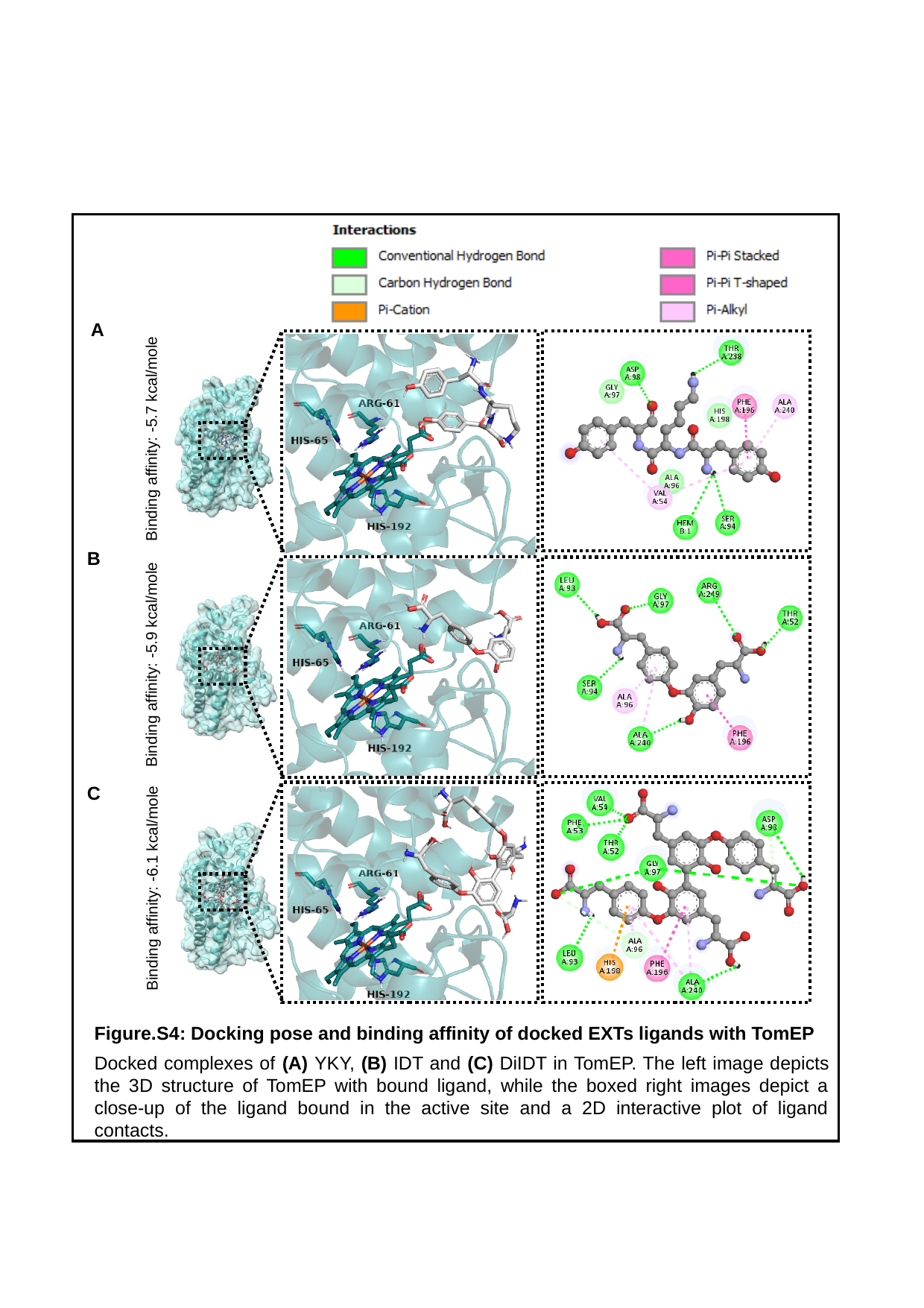

A
Binding affinity: -5.7 kcal/mole
B
Binding affinity: -5.9 kcal/mole
C
Binding affinity: -6.1 kcal/mole
Figure.S4: Docking pose and binding affinity of docked EXTs ligands with TomEP
Docked complexes of (A) YKY, (B) IDT and (C) DiIDT in TomEP. The left image depicts the 3D structure of TomEP with bound ligand, while the boxed right images depict a close-up of the ligand bound in the active site and a 2D interactive plot of ligand contacts.

### Slide 8
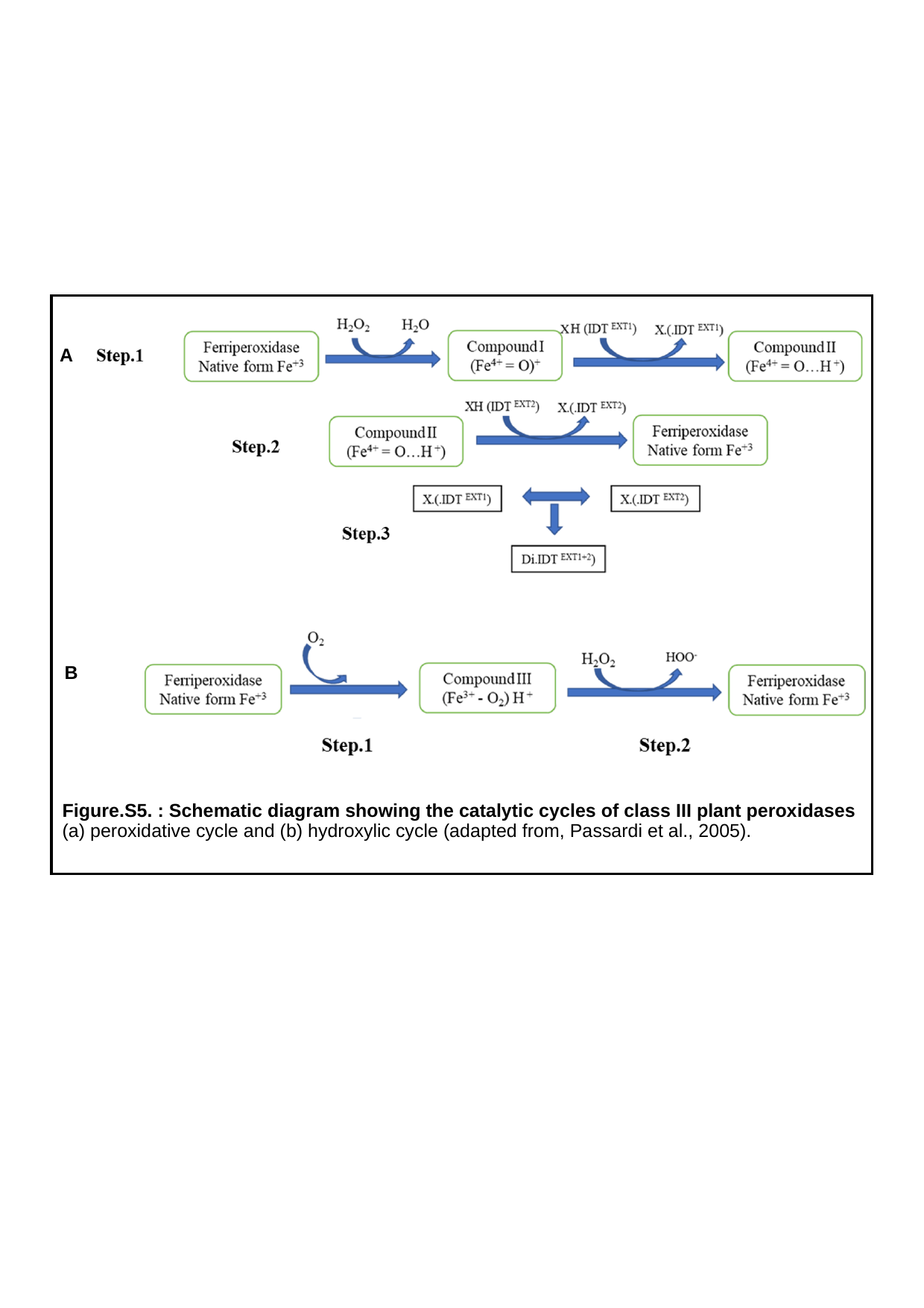

A
B
Figure.S5. : Schematic diagram showing the catalytic cycles of class III plant peroxidases
(a) peroxidative cycle and (b) hydroxylic cycle (adapted from, Passardi et al., 2005).
